# Classifying Genetic Interactions Using an HIV Experimental Study

**DOI:** 10.1101/2024.05.13.594050

**Authors:** Sean C. Huckleberry, Mary S. Silva, Jeffrey A. Drocco

**Affiliations:** Electrical Engineering and Computer Science, Massachusetts Institute of Technology, Cambridge, MA, USA; Global Security-Computing Applications, Lawrence Livermore National Labs, Livermore, CA, USA; Physical Life Sciences Directorate, Lawrence Livermore National Labs, Livermore, CA, USA

**Keywords:** Gene interaction, bioinformatics, machine learning

## Abstract

Current methods of addressing novel viruses remain predominantly reactive and reliant on empirical strategies. To develop more proactive methodologies for the early identification and treatment of diseases caused by viruses like HIV and Sars-CoV-2, we focus on host targeting, which requires identifying and altering human genetic host factors that are crucial to the life cycle of these viruses. To this end, we present three classification models to pinpoint host genes of interest. For each one, we thoroughly analyze the current predictive accuracy, susceptibility to modifications of the input space, and potential for further optimization. Our methods rely on the exploration of different gene representations, including graph-based embeddings and large foundation transformer models, to establish a set of baseline classification models. Subsequently, we introduce an order-invariant Siamese neural network that exhibits more robust pattern recognition with sparse datasets while ensuring that the representation does not capture unwanted patterns, such as the directional relationship of genetic interactions. Through these models, we generate biological features that predict pairwise gene interactions, with the intention of extrapolating this proactive therapeutic approach to other virus families.

## I. Introduction

In the ongoing pursuit of effective antiviral strategies, recent headway has been made in developing proactive therapies to improve public health by preventing or reducing the severity of infections and mitigating transmission among vulnerable groups. One main challenge, however, is identifying a subset of promising genes for proactive host targeting, as comprehensive therapy research is time- and cost-intensive. Once identified, groups of genes could provide a foundation for efficient validation studies and clinical trials. Crucially, these tasks could offer insight into common viral pathways that may be targeted to disrupt viral infection, which may be relevant to a broader group of viruses of interest. Here, we employ three predictive models and compare their efficacy in addressing this obstacle with a focus on Human Immunodeficiency Virus (HIV). Our interest in HIV stems from its well-characterized genetic mechanisms and abundant genetic data, which provide a robust framework for probing pertinent viral-host interactions.

Our models rely heavily on genetic pairwise epistasis: the interactions between pairs of genes. To model epistasis, we introduce two main featurization approaches. The first is based on graphical genetic relationships between genes, biological processes, pathways, and cellular components. These graph relationships are cached in the Scalable Precision Medicine Oriented Knowledge Engine (SPOKE), a “database of databases” comprised of approximately 20 thousand human genes and over 1 million gene expression and regulation edge types [1]. The second method uses Geneformer, a foundation transformer model pre-trained on 30 million single-cell transcriptomes [2]. We utilize the embedding outputs from this pre-trained model and perform fine-tuning with a task-specific neural network classifier to predict genetic epistasis. Using these featurization approaches, we present in the form of models a technique for identifying vital genes for host targeting, which we anticipate may be adapted for the treatment of other viruses beyond HIV.

## II. Methods

### A. Featurization Approaches

#### 1) Gene representations using graph embeddings

Our graph-based featurization approach employs the University of San Francisco’s Scalable Precision Medicine Oriented Knowledge Engine (SPOKE), a graphical network of biomedical databases that has been used for drug repurposing [3], gene regulation in lung cells for COVID-19 patients [4], and numerous other link prediction tasks. This knowledge graph consists of over 20 thousand protein-coding human gene nodes from Entrez Gene (NCBI’s database for gene-specific information) [5], a combined 18 thousand biological process, molecular function, and cellular component node types derived from the Gene Ontology database [7]. These nodes contain comprehensive gene information, including gene nomenclature, function and attributes, sequences, and source data. Additionally, the graph includes over 1 million gene relationships, where the knockdown or knockout of one gene, achieved either by short hairpin RNA or CRISPR-Cas9, results in the upregulation or downregulation of another gene as indicated by consensus transcriptional profiles. Using various subsets of nodes, we apply the Fast Random Projection (FastRP) graph embedding, a sparse random projection-based graph embedding algorithm [8]. With FastRP, we represent each gene node in the SPOKE graph as a 256-dimensional numeric vector. After extracting the embeddings associated with the 356 genes associated with HIV function (which match our validation set discussed below), we use the concatenated form as inputs to model corresponding gene-pair epistasis. This allows us to explore the relationship between gene embedding pairs and their corresponding interactions.

#### 2) Gene representations using Geneformer embeddings

Pretrained on the massive Genecorpus-30M, which consists of 30 million single-cell transcriptomes, Geneformer contains comprehensive information in its embeddings that opens new avenues for computation. Our second featurization approach involves employing these embeddings to perform inference regarding their impact on the epistatic gene interactions that contribute to HIV.

In the standard Geneformer architecture, each single-cell transcriptome is assigned a rank value encoding: genes are ranked by their expression in each cell normalized by their expression across the entire Genecorpus. This encoding system prioritizes genes that differentiate cell state and shrinks housekeeping genes with limited distinguishing potential [2]. The rank value genes are then passed into a six-layer BERT transformer encoding unit [9], and the model is pre-trained using a context-aware masked learning objective (Figure 1). We utilize the pre-trained model by extracting the final hidden layers of the ranked genes, which are 256-dimensional embeddings, and using them to compare both featurization methods.

**Fig. 1.**
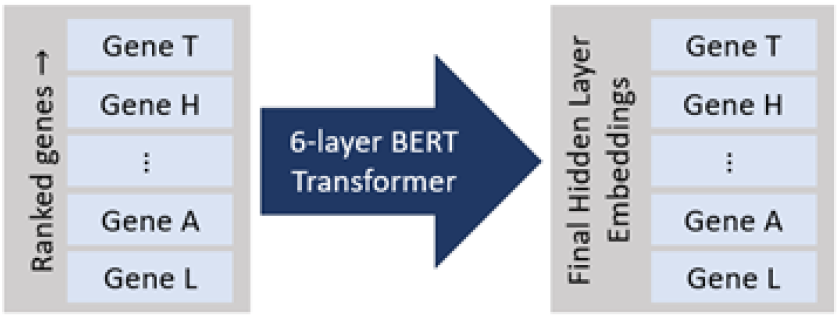
Geneformer architecture and feature extraction. Each layer within the Geneformer BERT Transformer has four attention heads that individually learn to monitor distinct classes of genes and improve predictive power.

We utilize the pretrained model by extracting the final hidden layers of the ranked genes, which are 256-dimensional embeddings, and use them to compare both featurization methods.

### B. Model Validation

As our predictive models harness the HIV infection metric as the target variable, an epistasis mapping containing 63,012 pairwise interactions closely linked to HIV function serves for validation. Generated as part of a recent study concerning the quantitative mapping of genetic interactions, an HIV epistasis matrix known as a vE-MAP (viral epistatic miniarray profile) was produced by the pairwise depletion of 356 human genes closely linked with HIV. More specifically, this vE-MAP approach involves the construction of miniarrays containing HIV and uniquely perturbed culture cells, and the resulting degree of infection of each cultured cell was captured and documented in the 356 by 356 symmetric epistatic matrix [**?**].

We focus on the upper triangular entries of the symmetric epistasis matrix, which serve as the response variables for our machine learning models. In scenarios where our objective is solely to classify HIV enhancement or suppression, we establish a threshold and formulate the response as a binary outcome.

## III. Results and Discussion

For each gene pair, the results of our predictive models are formatted as a binary response indicating HIV suppression or enhancement. To this end, we determine an appropriate threshold for the HIV infection metric and categorize response into two groups. Establishing the threshold as the mean of the response, we achieve a balanced split between suppression and enhancement, mitigating concerns related to imbalance (Figure 2). Of particular interest are gene pairs associated with HIV suppression, which may be crucial to developing host-targeting therapies.

**Fig. 2.**
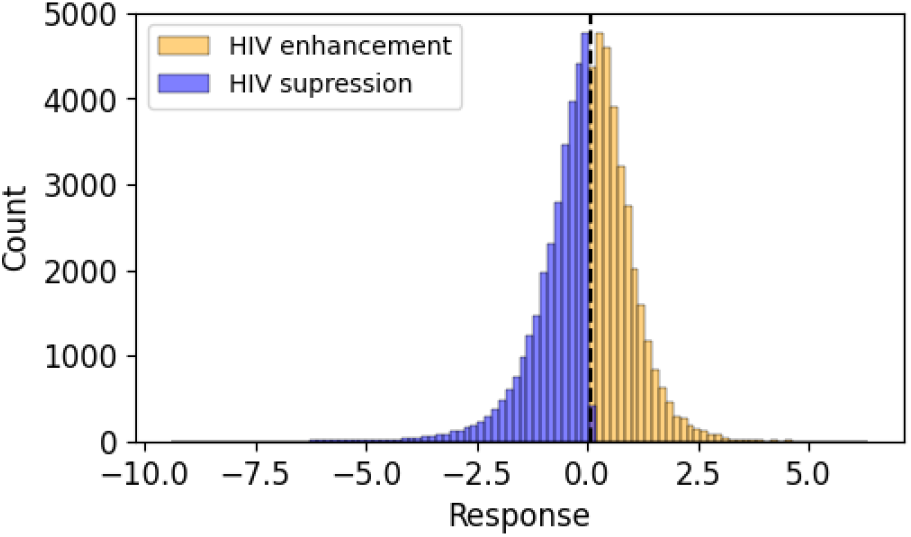
A histogram representing the epistasis between each pair of genes in the 356 by 356 matrix. The tails represent HIV enhancement on the positive end and suppression at the negative end. Most pairs of genes show no epistasis response at the center of the distribution. The red line indicates the threshold used for binary classification.

### A. FastRP Embedding Classification

The FastRP graph-based approach enables us to capture a different feature space from the Geneformer embeddings by focusing on a subset of genes and their relationships within the larger SPOKE knowledge graph. Utilizing the SPOKE and FastRP embeddings, we explore the “gene” node types, which encompass approximately 20 thousand human genes and contain gene names, descriptions, abbreviations, and source information. Similarly, we investigate the “biological process” node types, which consist of roughly 12 thousand nodes defining biological objectives to which a gene or gene product contributes. There is a discrete, fixed number of these nodes defined within Gene ontology, so the relationships between these node types and the genes are unique and finite. Figure 3 illustrates how we utilize each node type to predict a response.

**Fig. 3.**
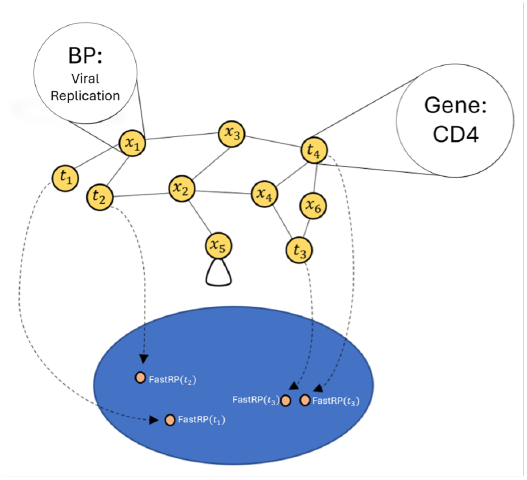
Graph-like representation of the data utilized by the FastRP predictive model. Target nodes *t*_*i*_ represent each of the 356 “gene” nodes from our validation set. Our model seeks to capture the latent representation of these target nodes using fast random projection embeddings FastRP(*t*_*i*_) and information regarding the neighbors in “biological process” nodes *x*_*j*_.

We begin by defining the binary response threshold, then shuffling and splitting the data so that 70% is used for training and 30% is reserved for validation. Subsequently, a random forest classifier was trained on the data to establish a predictive model, given the method’s exceptionally robust performance for datasets with many features [11]. The model was then evaluated on the test set to assess its performance and generalization capabilities.

This first model, which harnesses the concatenated FastRP embeddings and binarized epistatic response, obtains a prediction accuracy of approximately 70% (Table I). Results falling along the off-diagonals represent incorrect predictions about gene pair epistasis. We seek to minimize the false positives, in which a gene pair is erroneously classified as exhibiting HIV suppression despite actually promoting HIV enhancement. These data points constitute a subset of gene pairs whose presence would likely be damaging rather than beneficial to a host, and we strive to exclude them from the output of our models.

**TABLE I.** FastRP Embeddings Random Forest Model Confusion matrix representing the performance of random forest classifier trained on FastRP embeddings.

| Predicted | Actual |  |
| --- | --- | --- |
|  | Enhancement | Suppression |
| Enhancement | 36.01% | 13.99% |
| Suppression | 15.92% | 34.08% |

### B. Geneformer-based Classification

We now explore the Geneformer embeddings using the same train-test splits and Random Forest parameters from the FastRP model. Notably, the Geneformer embeddings under analysis were derived from the final layer, which is recognized for capturing features more closely associated with the learning objective of the pre-trained task. Despite the lack of fine-tuning, both featurization approaches lead to fair prediction accuracy.

The prediction accuracy of the Geneformer and FastRP models is nearly identical (Table II). We conclude that both featurization approaches offer similar predictive performance as the training and testing parameters were kept constant.

**TABLE II.** Geneformer Embeddings Random Forest Model Confusion matrix representing the performance of random forest classifier trained on Geneformer embeddings.

| Predicted | Actual |  |
| --- | --- | --- |
|  | Enhancement | Suppression |
| Enhancement | 35.98% | 14.02% |
| Suppression | 15.87% | 34.13% |

This observation is confirmed by comparing the ROC/AUC of the two models, plotting true positive against false positive rates (Figure 4). Notably, Geneformer embeddings demonstrate only marginally superior performance compared to the FastRP embeddings.

**Fig. 4.**
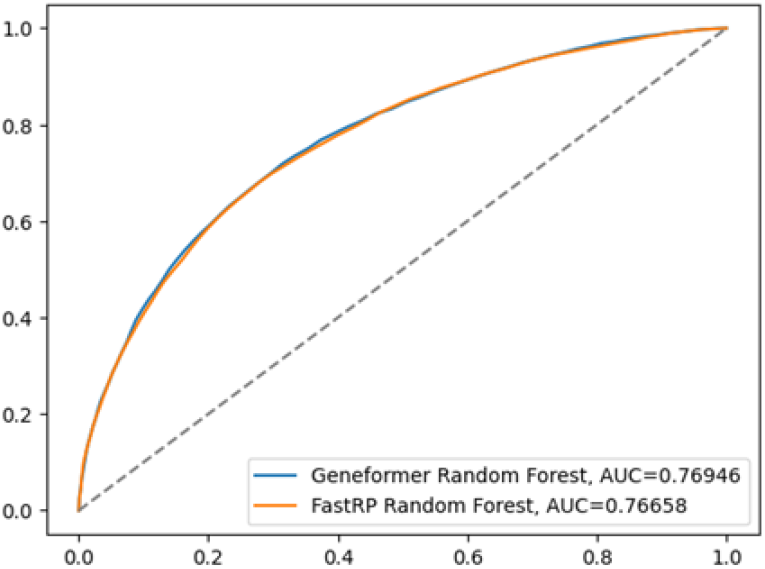
ROC/AUC curve comparing the predictive performance of the Geneformer and FastRP models. Random chance is represented by the dotted linThe dotted line represents random chance, and the “perfect classifier” reaches the top left corner of the graph. The overall performance of each model can be summarized as a scalar: the area under the curve (AUC).

#### 1) Symmetry and Order Invariance

Order invariance is vital for our epistasis modeling that utilizes a symmetric binary input of gene pairs since it indicates that any trends learned by the model reflect the biological interaction between genes and are not artifacts of changes in the data representation. However, both the FastRP embedding classification and Geneformer-based classification models ignore the order of embedding concatenation. In other words, [EmbeddingGeneA, EmbeddingGeneB] and [EmbeddingGeneB, EmbeddingGeneA] may yield inconsistent results despite being pairwise symmetrical. To demonstrate this, we compare the predicted results of the Geneformer embedding with the same training inputs and random forest parameters but swapped concatenation order of the observations in the test set. Swapping the concatenation order of gene pairs resulted in a 23% inconsistency between predictions (Table III). This reveals the existence of a disagreement in the prediction of the same observations when the concatenation order is swapped, indicating that the model has captured extraneous patterns during training. We address this concern by implementing a Siamese network.

**TABLE III.**
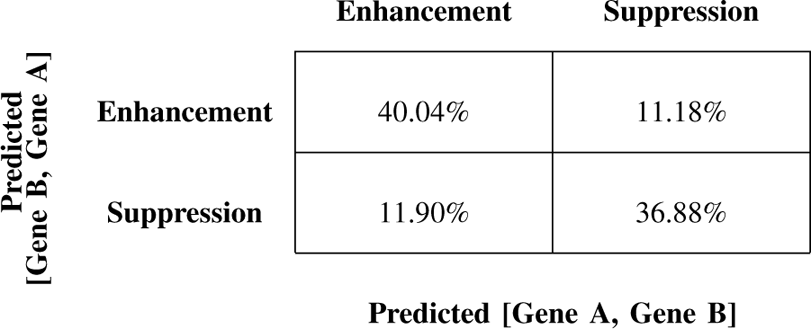
Geneformer Random Forest Permuted Feature Space Confusion matrix comparing the binarized output of symmetric inputs for random forest classifier trained on Geneformer embeddings. For an order-invariant model, one would expect that the values of the off-diagonals are close to zero.

### C. Siamese Classification Network

We implement a Siamese network to ensure cohesion between gene pairs (Figure 5). This type of network better correlates symmetric gene pairs and introduces order-invariance to our model. Siamese neural networks have been studied in the context of machine learning for classifying similarities or dissimilarities between image inputs as well as signature verification [12]. They have also been used for protein-protein interaction classification problems where the input space is the embeddings extracted from protein sequence transformer models, commonly referred to as ProtBERT [13]. Since we determined that FastRP and Geneformer produce similar results, we focus on the Geneformer embeddings to implement this network.

**Fig. 5.**
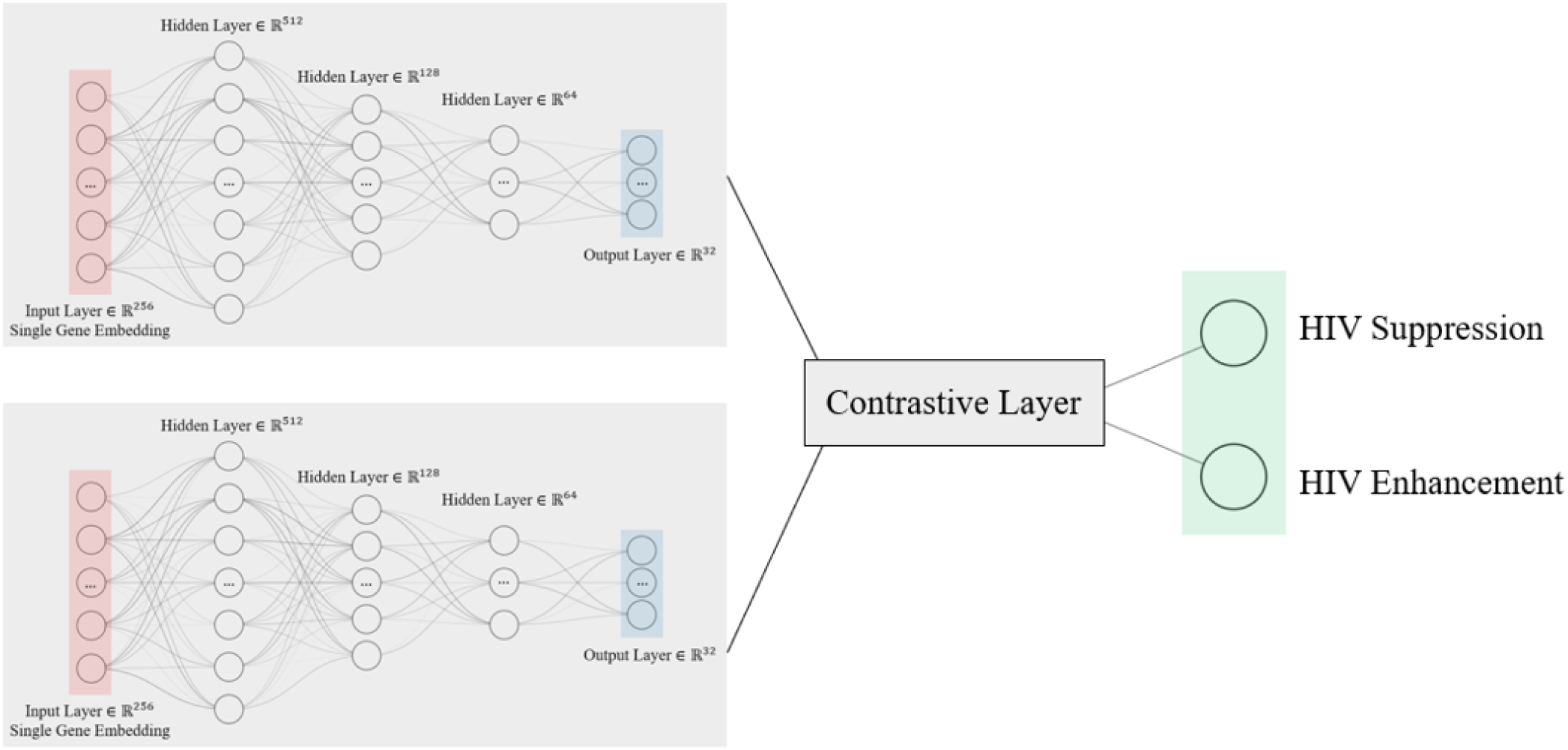
Siamese network architecture. The model includes two identical neural networks that are applied independently to each gene embedding. The output layers of the two networks are then fed into a contrastive layer which produces a single output containing information about the final similarity or dissimilarity between the input pairs.

The Siamese network is significantly more sensitive to hyperparameter tuning than the random forest models as it consists of two identical neural networks, where the inputs are the corresponding gene embeddings. Much like a standard feed-forward neural network, the number of hidden layers and corresponding weights and biases, and the choice of the optimizer, activation functions, and regularization techniques can significantly impact the performance and accuracy of a Siamese neural network. Initially, we define the two branches of our Siamese network to have three hidden layers of size 512, 128, and 64. We select a stochastic gradient descent optimizer, which is memory-efficient and computationally inexpensive relative to other optimizers. The contrastive layer, used to measure the similarity of inputs, is defined as a concatenation of the layers of the two Siamese network branches and their Euclidean distance with an additional linear layer. Other contrastive layers to explore include a function of the Euclidean distance or a function of the cosine similarity. Due to the large scale of the data inputs and complexity of the Siamese neural network, mini-batch training and PyTorch GPU acceleration were leveraged.

Beyond improving the model’s consistency for symmetric gene pairs, the Siamese network, trained on the Geneformer embeddings, also slightly enhanced the model’s predictive accuracy from approximately 70% to over 71% without additional fine-tuning (Table IV). In contrast with the previous two models, which have limited improvement potential from the current state, this Siamese network could see a vast improvement in predictive accuracy from tweaking the hyperparameters, either manually or with an optimization framework like Optuna.

**TABLE IV.** Geneformer Embeddings + Siamese Neural Network Confusion matrix comparing the binarized output of symmetric inputs for a Siamese network trained on Geneformer embeddings.

| Actual |  |  |
| --- | --- | --- |
|  | Enhancement | Suppression |
|  | Predicted |  |
| Enhancement | 35.41% | 14.59% |
| Suppression | 14% | 36% |

The main benefit of applying a Siamese Neural Network to the Geneformer embeddings is that it solves the problem of feature input order invariance. Unlike the first model, which saw approximately 23% disagreement of the test set prediction when the order of the gene embeddings was swapped, the Siamese Neural Network now has 100% agreement (Table V). This also indicates that the model is not unintentionally learning patterns associated with the order of the inputs.

**TABLE V.** Geneformer Embeddings + Siamese Neural Network Permuted Feature Space Confusion matrix comparing binarized output of symmetric inputs for Siamese network trained on Geneformer embeddings. The off-diagonals are zero, as expected for an order-invariant model.

| Predicted<br>[Gene B, Gene A] |  |  |
| --- | --- | --- |
|  | Enhancement | Suppression |
|  | Predicted [Gene A, Gene B] |  |
| Enhancement | 49.41% | 0% |
| Suppression | 0% | 50.59% |

## IV. Conclusion

To better understand host-targeting viral therapies, three models were trained to classify gene pairs as inducing HIV suppression or enhancement. The first classification model explored the FastRP embeddings from UCSF’s SPOKE database with a graph-based approach and achieved a prediction accuracy of 70% with a random forest algorithm. The next model, which achieved similar performance, utilized the same algorithm and parameters, but embeddings were extracted from the pre-trained Genformer foundation transformer instead. The first pair of models employ a random forest classifier, utilizing the FastRP and Geneformer embeddings as individual inputs without undergoing fine-tuning. These models perform adequately when trained on a sizable subset of data. However, the potential for significant improvement is limited due to the inherent characteristics of random forests. Random forests rely on decision trees, which independently make decisions based on individual features and do not provide much flexibility in capturing non-linear relationships and other hierarchical relationships between the input space. More importantly, with the concatenated representation of the input feature space, they fail to capture the symmetric relationship between genetic interaction.

The third model employed a Siamese network to the Geneformer embeddings, which introduced order-invariance to better handle input symmetry and demonstrated significant optimization potential through fine-tuning. By inferring gene pairs inducing HIV suppression, these models identify the most promising pairs for HIV therapies to study through more resource-intensive methods. The next steps include applying these models to different diseases and sparse datasets where they might be most informative. Regarding genetic interaction, we have only explored these models under a binary classification setting; a stricter multi-class response could include a third neutral class. The overall accuracy may be reduced, but using a multi-class response would give a more realistic identification of genetic interactions, as it would produce more nuanced results.

This experimental study underscores the critical role of various computational models in adeptly predicting HIV’s associated human host factors through rigorous pattern matching of large-scale gene data. The significance of this work is manifold: it further establishes data and computation as a method to expedite and reduce the cost of traditional research methods in biology, and suggests that predictive models can enable us to take a more proactive approach to understanding viral infections.

## Acknowledgments

This work was performed under the auspices of the U.S. Department of Energy by Lawrence Livermore National Laboratory under Contract DE-AC52-07NA27344 (LLNL-JRNL-860169) This document was prepared as an account of work sponsored by an agency of the United States government. Neither the United States government nor Lawrence Livermore National Security, LLC, nor any of their employees makes any warranty, expressed or implied, or assumes any legal liability or responsibility for the accuracy, completeness, or usefulness of any information, apparatus, product, or process disclosed, or represents that its use would not infringe privately owned rights. Reference herein to any specific commercial product, process, or service by trade name, trademark, manufacturer, or otherwise does not necessarily constitute or imply its endorsement, recommendation, or favoring by the United States government or Lawrence Livermore National Security, LLC. The views and opinions of authors expressed herein do not necessarily state or reflect those of the United States government or Lawrence Livermore National Security, LLC, and shall not be used for advertising or product endorsement purposes.

## Notes

### Competing Interest Statement

The authors have declared no competing interest.

